# Structural insights into human ABCC4 mediated transport of platelet agonist and inhibitor

**DOI:** 10.1101/2022.12.25.521909

**Authors:** Yu Chen, Liang Wang, Wen-Tao Hou, Zhi-Hui Zha, Kang Xu, Cong-Zhao Zhou, Qiong Li, Yuxing Chen

## Abstract

The activation and accumulation of human platelets contributes to hemostasis and thrombosis, the imbalance of which would cause cardiovascular diseases, an increasing threat to the global health. Human ABC transporter ABCC4 that pumps out the platelet agonist and anti-platelet drug such as aspirin, might become a promising target for preventing cardiovascular diseases. Here we solve the structures of human ABCC4 in the apo and two complexed forms, all of which adopt a typical architecture of type-IV ABC transporters in an inward-facing conformation. Structure of ABCC4 complexed with U46619, an analog of the unstable TXA_2_, provides the first structural evidence that the platelet agonist TXA_2_ is also exported via ABCC4. The dipyridamole-complexed structure reveals the inhibitory mechanism of dipyridamole against ABCC4. Structural comparisons enabled us to identify a transmembrane pocket in ABCC4 that provides a defined space for the rational design of specific anti-platelet drugs.

## Main

ATP-binding cassette subfamily C contains 12 members in total, all of which adopt a canonical architecture of ABC exporters with two nucleotide-binding domains (NBDs) and two transmembrane domains (TMDs) of 6+6 transmembrane helices^1^. Except for ABCC7 (the regulator of cystic fibrosis) and ABCC8/ABCC9 (the receptors of sulfonylurea), nine members of ABCC transporters, namely ABCC1~ABCC6 and ABCC10~ABCC12, were first discovered and well-known as multidrug resistance proteins (MRPs), sequentially termed MRP1~MRP9^2^ These MRPs could extrude various drugs out of the cell in an ATP-dependent manner, thus leading to drug resistance in chemotherapy of cancer and antiviral/bacterial treatment^3^. Moreover, they are also responsible for the secretion of a wide spectrum of physiological substrates in various cellular processes^4–8^, such as cAMP, glutathione (GSH), leukotriene C_4_ (LTC_4_), bile acids and steroid hormone.

ABCC4/MRP4, encoded by *ABCC4* gene located on chromosome 13q32.1 of the human genome^9^, possesses three splicing variants, isoform 1 of which is the full-length version and well-known canonical ABCC4. Previous reports showed that it is widely expressed in most human tissues, including the platelet, prostate, heart, liver and intestine^10,11^. ABCC4 was first identified as a homolog of ABCC1/MRP1 and ABCC2/MRP2^9^, and proved to transport a wide range of clinic drugs, especially the cardiovascular, antiviral, antibiotic and cytotoxic agents^11–17^. Moreover, besides the exogenous substrates, ABCC4 is also capable of pumping a variety of physiological substrates out of the cell in various tissues.

It was reported that ABCC4 is highly expressed in human platelets, and subsequently presents on the platelet membrane and in dense granules^18^. The in vitro assays proved that ABCC4 could transport thromboxane TXB_2_^19^, a stable metabolite derived from the platelet agonist TXA_2_^19–22^. As we know, the extracellular TXA_2_ is secreted from the platelets, and binds to the thromboxane-prostanoid-α receptor on the platelet membrane, eventually induces the platelet activation and accumulation^23^. The activated platelets could further recruit platelets from the flowing blood, contributing to hemostasis upon vascular injury, but leading to thrombosis of cardiovascular diseases under pathophysiological conditions. Despite TXA_2_ is a poorly membrane permeable molecule that could be partly diffused from the platelet^24^, an active exporter for TXA_2_ was hypothesized to be necessary for its instant and effective secretion.

Export of the agonist ADP from the platelet via ABCC4 is also crucial to maintain the function of platelets^18^. The ADP analog dipyridamole, which is used in clinic as a coronary vasodilator, could specifically inhibit ABCC4 and attenuate the efflux of ADP from the platelets, thus preventing the thrombus formation^18^. In addition, ABCC4 could also pump out aspirin, a widely used anti-platelet drug in clinic for the inhibition of thromboxane synthesis^25^. In some clinical cases, such as the secondary stroke prevention and treatment of coronary heart disease in patients with noncardioembolic ischemic stroke or transient ischemic attack, coadministration of dipyridamole with low-dose aspirin is recommended^26^.

Altogether, ABCC4 might become a promising target of anti-platelet drugs for cardiovascular diseases. However, its fine structure and transport mechanism towards such a variety of substrates remain unknown. Here, using single-particle cryogenic electron microscopy (cryo-EM), we solved the three-dimensional (3-D) structures of human ABCC4 in the apo form and complexed forms bound to physiological substrate/clinical inhibitor. Structural analyses in combination with biochemical assays give evidence that the platelet agonist TXA_2_ is the substrate of ABCC4, and reveal a specific transmembrane substrate-binding pocket of ABCC4. These findings will shed light on the development of new drugs against the cardiovascular diseases.

## Results

### The ATPase activities of ABCC4 upon the addition of substrate or inhibitor

The full-length human ABCC4 of 1325 residues was recombinantly overexpressed in human embryonic kidney (HEK) 293F cells, extracted from the membrane, and purified with n-dodecyl-β-d-maltoside (DDM) plus cholesteryl hemisuccinate (CHS). To obtain the stable and homogeneous proteins for structure determination and the mimetic physiological membrane environment for biochemical assays, we further reconstituted the detergent-solubilized ABCC4 into lipid nanodiscs (Extended Data Fig. 1). The purified ABCC4 possesses a basal ATPase activity at 18.07 mol Pi min^-1^ mol^-1^ protein, whereas the double mutant ABCC4^D560N&E1202Q^ (the degenerated aspartate and catalytic glutamate in Walker B motifs of NBD1 and NBD2 replaced with asparagine and glutamine, respectively) displayed a great decrease of ATP hydrolysis activity (Fig. 1a).

**Fig. 1.**
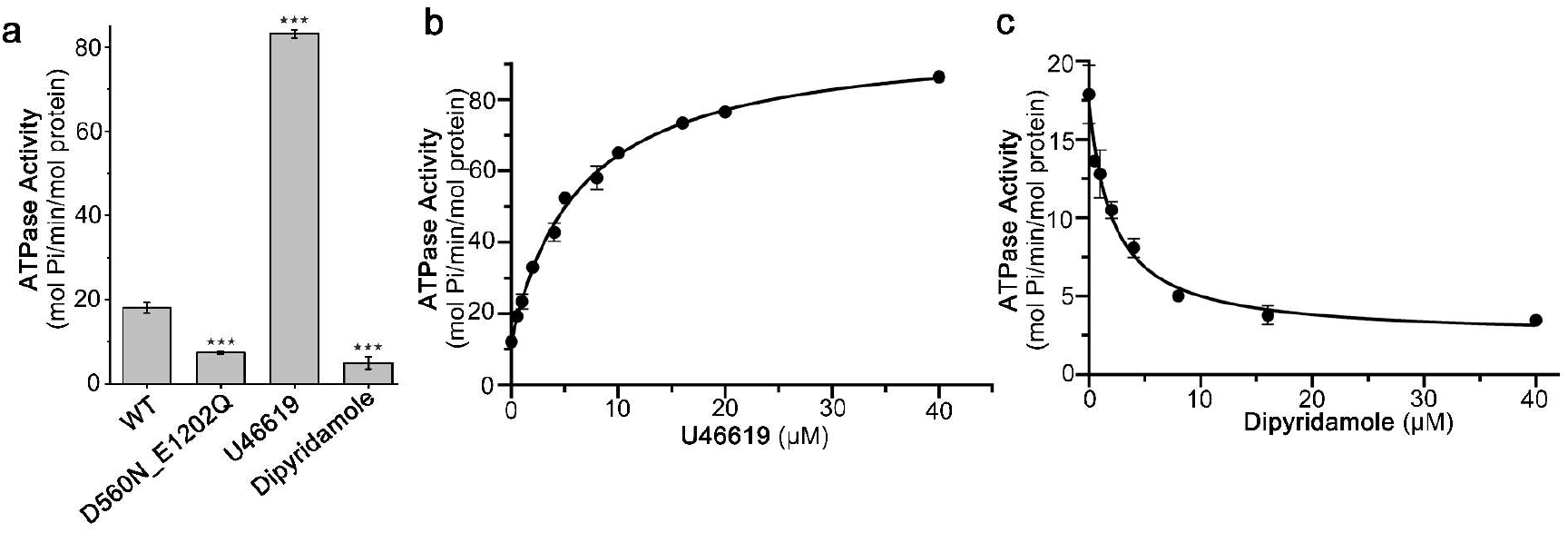
Substrate-stimulated/inhibited ATPase activity assays of ABCC4. **a** The ATPase activities of wild-type (wt) ABCC4, the catalytic mutant ABCC4^D560N&E1202Q^, and ABCC4 in the presence of substrate U46619 and inhibitor dipyridamole.**b-c** Substrate concentration-dependent ATPase activity of ABCC4 in the presence of 2 mM ATP upon addition of **b** U46619 and **c** dipyridamole at the increased concentrations. The data points are fitted with Logistic equation. All data points are the average of three independent experiments, and error bars represent the means ± SD. The protein is purified in nanodiscs.

As reported, ABCC4 could export PGE_2_^22^, a most abundant prostaglandin that participates in multiple physiological pathways, such as immune response, female reproduction and cancer progression^27,28^. Considering that TXA_2_ shares a chemical structure strongly similar to PGE2 (Extended Data Fig. 2), we proposed that TXA_2_ might also be actively exported by ABCC4, which is localized on the platelet membrane^18^. As TXA_2_ is prone to degradation under aqueous conditions due to an unstable ether linkage, U46619, a stable analog of TXA_2_^29^, was applied to the ATPase activity assays. As predicted, addition of U46619 triggers a 4-fold increase of ATPase activity, in comparison with the basal ATPase activity of apo-form ABCC4 (Fig. 1a). Moreover, U46619 possesses a half maximal effective concentration (*EC*_50_) of ~6.4 μM and *V*_max_ of ~83.1 mol Pi min^-1^ mol^-1^ protein (Fig. 1b), indicating a high in vitro affinity with ABCC4.

In contrast to U46619, the biochemical assays showed that the ATPase activity of ABCC4 is largely attenuated upon the addition of dipyridamole (Fig. 1a). Furthermore, the concentration-dependent activity curve of dipyridamole yields a median inhibitory concentration (*IC*_50_) of ~2.1 μM and *V*_min_ of ~2.4 mol Pi min^-1^ mol^-1^ protein, respectively (Fig. 1c). It suggested that ABCC4 has a slightly higher binding-affinity towards dipyridamole, compared to U46619.

### Overall structure of apo-form ABCC4

To obtain its 3-D structure, the purified ABCC4 in lipid nanodiscs was applied to cryo-EM sample preparation and data collection (Extended Data Fig. 3 a, b). Using 392,047 particles in the best class, the structure of apo-form ABCC4 was solved at an overall resolution of 3.13 Å based on the gold-standard Fourier shell correlation (FSC) 0.143 criterion (Extended Data Fig. 3c-f). In total, 1232 residues were *de novo* built according to the EM density, except for the N-terminal 8 and C-terminal 27 residues, in addition to a long linker (residues Asp634-Val696) between NBD1 and TMD2.

ABCC4 is a full transporter that consists of two TMDs and two NBDs in one polypeptide, displaying as a pseudo-symmetric homodimer (Fig. 2a). Its apo-form structure adopts an inward-facing conformation (Fig. 2a, b), similar to the previously reported type-IV apo-form ABC homologs, such as human ABCB11 and ABCD1^30,31^. Each TMD of ABCC4 is composed of six transmembrane helices, a significant portion of which extends to the cytosol, leading to the NBDs protruded far away from the membrane (Fig. 2a). The TMDs form two diverged “wings” at the intracellular side, but tightly pack against each other at the outer membrane leaflet (Fig. 2a). Notably, helices TM4 and TM5 from TMD1, together with TM10 and TM11 from TMD2, are domain-swapped to the opposite TMD (Fig. 2a), which is a key feature of type-IV ABC transporters^1^. The NBDs of ABCC4 possess a canonical NBD fold of ABC transporters (Fig. 2a), in which the Walker B motif of NBD1 has a degenerated active site, namely that the catalytic residue glutamate is replaced by aspartate. Moreover, two pairs of coupling helices from the TMDs are embedded in the grooves on the NBDs (Fig. 2a) with a buried interface area of 2158 Å^2^. In addition, the lasso motif at the N-terminus, and two classic elbow helices, are finely modeled in the apo-form ABCC4 structure.

**Fig. 2.**
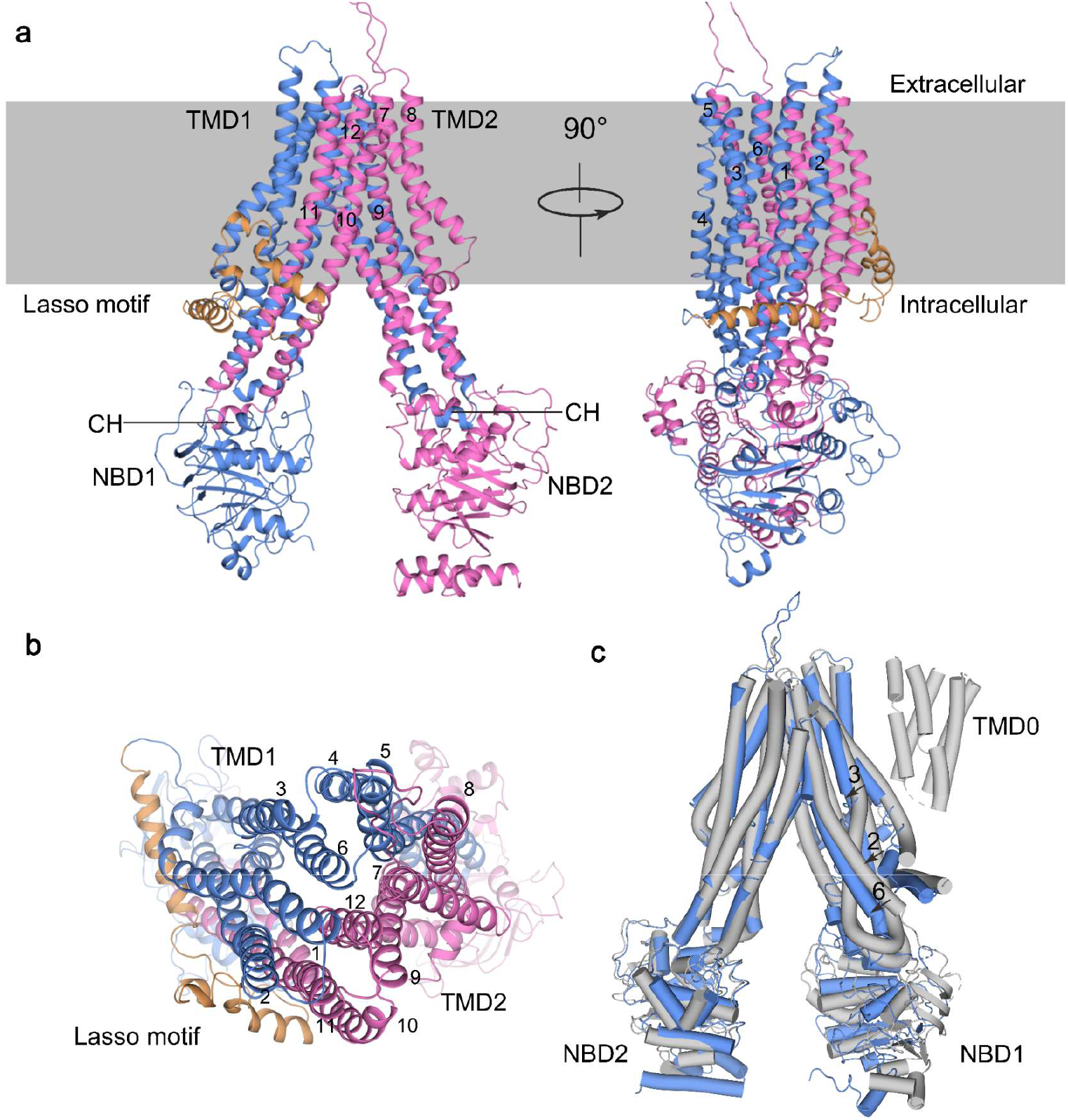
Overall structure of apo-form ABCC4. **a** Cartoon representation of the apo-form ABCC4. The two halves of ABCC4 are colored in blue and magenta for half 1 (TMD1 and NBD1) and half 2 (TMD2 and NBD2), respectively. The N-terminal lasso motif is colored in orange. The coupling helices (CH) are indicated. Transmembrane helices (TMs) are labeled. The membrane plane is indicated as the gray rectangle. **b** Top view of the apo-form ABCC4. The 12 TMs are sequentially numbered. **c** Superposition of the apo-form human ABCC4 (blue) against the apo-form bovine MRP1 (bMRP1, gray, PDB code: 5UJ9). The TMs that possess conformational changes are labeled. The additional TMD0 of bMRP1 and the NBDs are indicated.

To date, among the nine MRPs, only the structure of bovine ABCC1/MRP1 (bMRP1) has been reported^6,32^. Compared to our apo-form structure, the apo-form bMRP1 also adopts an inward-facing conformation, but with an additional five-TM motif termed TMD0 at the most N-terminus (Fig. 2c). Superposition of these two apo-form structures yielded a root-mean-square-deviation (RMSD) of 2.7 Å over 912 aligned Cα atoms. Except for the two not-well-aligned NBD1, the intracellular parts of helices TM2, TM3 and TM6 of TMD1 in ABCC4 slightly approach towards TMD2, compared to those in bMPR1 (Fig. 2c).

### Structure of U46619-bound form indicates TXA_2_ is a putative physiological substrate of ABCC4

To reveal the substrate-binding pattern, U46619 was incubated with purified ABCC4 in lipid nanodiscs, and then applied to cryo-EM structure determination. Eventually, the 3-D structure of human ABCC4 complexed with U46619 was obtained at the resolution of 3.08 Å, in which the EM density of one U46619 molecule could be clearly observed in the transmembrane cavity (Extended Data Fig. 4). In general, the U46619-bound ABCC4 structure adopts an inward-facing conformation quite similar to the apo-form structure (Fig. 3a, b); and superposition of the two structures resulted in an RMSD of 1.102 Å over 1207 Cα atoms (Extended Data Fig. 5a), suggesting that no global conformational changes occurred upon U46619 binding.

**Fig. 3.**
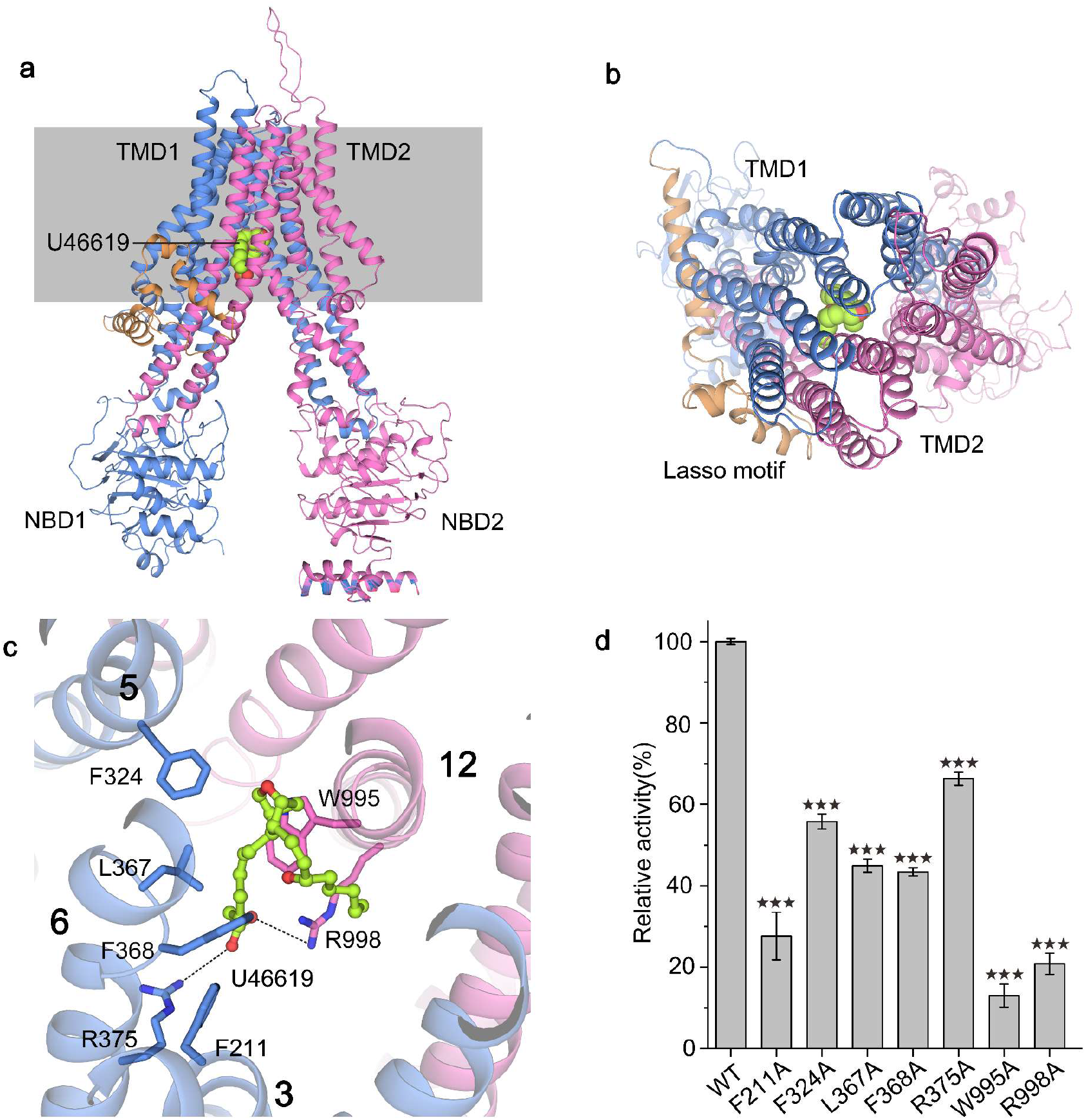
Overall structure of ABCC4 in complex with substrate U46619 and the substrate-binding pocket. **a-b** Cartoon representation of the U46619-bound ABCC4 in **a** side view and **b** top view. The two halves of ABCC4 are colored in blue and magenta for half 1 (TMD1 and NBD1) and half 2 (TMD2 and NBD2), respectively. The N-terminal lasso motif is colored in orange. The U46619 molecule is shown as limon spheres. The membrane plane is indicated as the gray rectangle. **c** Close-up view of the U46619-binding pocket. U46619 is shown as stick spheres in limon. The binding residues are shown as sticks and labeled. The hydrogen bonds are indicated by black dotted lines. **d** Relative ATPase activities of ABCC4 and mutants in the presence of 2 mM ATP upon addition of U46619. The relative activity represents the substrate-stimulated activity of ABCC4 or the mutants harboring a single mutation of residue at the substrate-binding pocket. Standard deviation is used for the comparison of statistical significance of mutants and wt. The *p* values of <0.05, 0.01, and 0.001 are indicated with *, ** and ***.

Structural analysis revealed that U46619 is bound to a pocket in the membrane (Fig. 3a, b), at a depth similar to that of LTC_4_ in bMRP1 (Extended Data Fig. 5b). This overall hydrophobic pocket is formed by residues Phe211 from TM3, Phe324 from TM5, Leu367 and Phe368 from TM6, and Trp995 from TM12, in addition to two hydrophilic residues Arg375 from TM6 and Arg998 from TM12 that contribute two pairs of hydrogen bonds (Fig. 3c). It is worth noting that, compared with the apo-form structure, residue Arg998 undergoes obvious induced fit upon substrate binding (Extended Data Fig. 5c).

Moreover, single-mutations of the interacting residues all lead to a significant decrease of the U46619-simulated ATPase activity of ABCC4 (Fig. 3d). In fact, multiple-sequence alignment revealed that most of these U46619-binding residues are highly conserved among ABCC4 homologs (Extended Data Fig. 6). Altogether, the specific binding of U46619 to ABCC4 is mainly maintained via extensive hydrophobic interactions, in addition to a hydrogen bond with the conserved polar residue Arg998. It is different from the reported bipartite LTC_4_-binding pocket of bMRP1, which contains two distinct moieties: a positively charged region and a largely hydrophobic area^6^. Considering that U46619 is an analog of TXA_2_, which should be secreted from the platelets, we proposed that TXA_2_ is most likely a physiological substrate of ABCC4.

### The structure of ABCC4 in complex with the inhibitor dipyridamole

The well-known inhibitor of ABCC4, dipyridamole (Extended Data Fig. 2), is a purine analog. It is widely used as an anti-platelet medicine in the clinical prevention or treatment of many diseases, including heart valve replacement, arterial grafting, cerebrovascular disorders, and disorders of peripheral circulation^33^. However, its inhibitory pattern against ABCC4 remains unknown. Using cryo-EM, we solved the 3.40 Å structure of ABCC4 in lipid nanodiscs pre-incubated with dipyridamole (Extended Data Fig. 7).

The overall structure of ABCC4 complexed with dipyridamole also adopts an inward-facing conformation (Fig. 4a), similar to that of apo- or complexed form ABCC4 (Fig. 2a and Fig. 3a). Superposition of dipyridamole-bound ABCC4 against apo-form and U46619-bound ABCC4 resulted in an RMSD of 0.697 Å over 1161 Cα atoms and 0.488 Å over 984 Cα atoms, respectively (Extended Data Fig. 8 a, b). Further structural analysis showed that the EM density in the transmembrane cavity is finely fitted with one dipyridamole molecule (Fig. 4a, b), which binds to a location almost the same to that of substrate U46619 (Extended Data Fig. 8b).

**Fig. 4.**
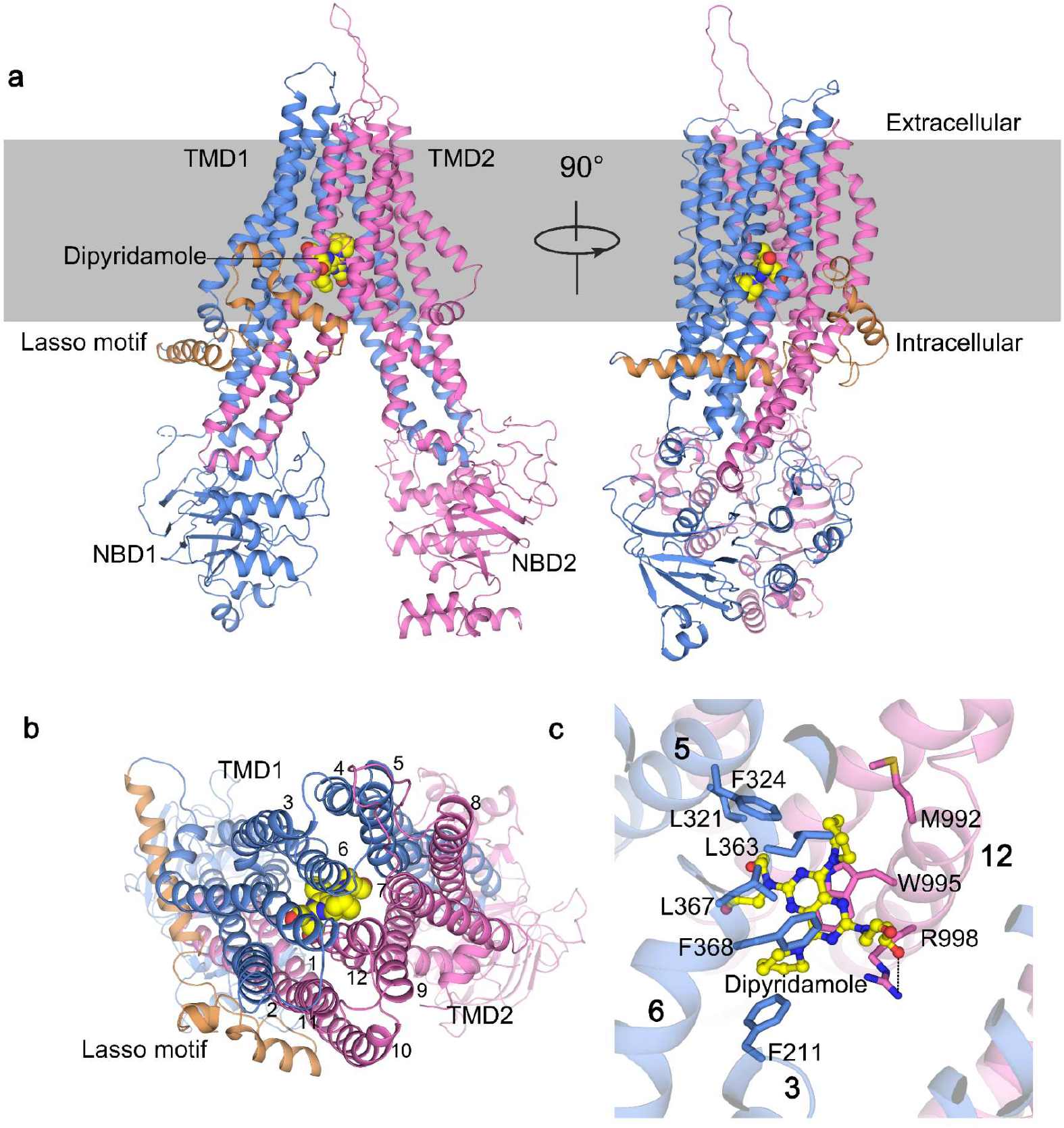
Overall structure of dipyridamole-bound ABCC4 and the inhibitor-binding pocket. **a** Cartoon representation of the dipyridamole-bound ABCC4. The two halves of ABCC4 are colored in blue and magenta for half 1 (TMD1 and NBD1) and half 2 (TMD2 and NBD2), respectively. The N-terminal lasso motif is colored in orange. The dipyridamole molecule is shown as yellow spheres. The membrane plane is indicated as the gray rectangle. **b** Top view of the dipyridamole-bound ABCC4. The TMs are numbered. **c** Close-up view of the dipyridamole-binding pocket. Dipyridamole is shown as stick spheres in yellow, and the binding residues are shown as sticks and labeled. The hydrogen bonds are indicated by black dotted lines.

Dipyridamole is bound to the substrate-binding pocket composed of a series of hydrophobic residues (Phe211 from TM3, Leu321 and Phe324 from TM5, Leu363, Leu367 and Leu368 from TM6, Met992 and Trp995 from TM12), in addition to Arg998 from TM12 via a hydrogen bond (Fig. 4c). Structural superposition against the structure of apo-form ABCC4 showed that no local conformational changes occurred upon binding of dipyridamole (Extended Data Fig. 8c). Notably, multiple-sequence alignment showed that except for the aforementioned substrate-binding residues, dipyridamole forms extra interactions with the residues Leu321, Leu363 and Met992 (Extended Data Fig. 6). It indicated that dipyridamole is a competitive inhibitor that binds to the same pocket of ABCC4 substrate, but at a higher affinity.

## Discussion

Human platelets regularly play critical roles in hemostasis; but their improper accumulation under pathophysiological conditions usually leads to a series of cardiovascular diseases, such as acute coronary syndrome, ischemic stroke and pulmonary embolism. As the potent platelet agonists, ADP and TXA_2_ stimulate the activation and accumulation of platelets via binding to receptors on the platelet membrane, and in turn induce hemostasis or thrombosis^23^. TXA_2_ also possesses vasoconstrictor property^27^, which increases the risk of cardiovascular diseases. To function at the extracellular side, it is prerequisite to export these two agonists out of the human platelets. Although ADP has been reported to be exported by the transporter ABCC4 that localized at the platelet membrane, it lacks direct evidence on how TXA_2_ is instantly transported out of platelets. Our ATPase activity assays towards a synthetic compound U46619, which perfectly mimic the chemical structure of the unstable TXA_2_, clearly showed that TXA_2_ is a specific substrate of ABCC4 (Fig. 1). Meanwhile, the structure of ABCC4 in complex with U46619 (Fig. 3) provided the first structural evidence that TXA_2_ is most likely pumped out of platelets via ABCC4.

Inhibition of the activation and/or accumulation of platelets is an effective and preferred approach for the anti-thrombotic treatment or the mitigation of disease progression in patients with cardiovascular diseases^34^. Considering that ABCC4 is responsible to the transport of two major platelet agonists, ADP and TXA_2_, competitive inhibitors of ABCC4 might become promising anti-thrombotic drugs. Actually, the ABCC4 inhibitor, dipyridamole, has been widely used as an anti-platelet drug in clinic ^18,26^. The present structure of ABCC4 in complex with dipyridamole clearly illustrated the dipyridamole-binding pocket (Fig. 4) and the inhibitory mechanism of dipyridamole against ABCC4.

Moreover, it was reported that high levels of ABCC4 proteins in platelets of patients after coronary artery bypass grafting surgery would lead to the resistance of aspirin, which is also a substrate of ABCC4 and usually used to prevent thrombosis and reduce cardiovascular complications in high-risk patients. In addition, ABCC4 was reported to regulate the cellular level of cAMP and control human leukemia cell proliferation and differentiation, making it also becomes a potential target for leukemia therapy^35^. Altogether, combined with previous reports, our findings indicated that ABCC4 is a promising target of anti-platelet drugs for cardiovascular and blood diseases; and give hints for the further design of clinical drugs to specifically inhibit the function of ABCC4.

## Acknowledgments

We thank Dr. Yongxiang Gao at the Center for Integrative Imaging, Hefei National Laboratory for Physical Sciences at the Microscale, University of Science and Technology of China during cryo-EM image acquisition for ABCC4. This work was supported by the Ministry of Science and Technology of China (2019YFA0508500) and The Strategic Priority Research Program of the Chinese Academy of Sciences (XDB37020202).

## Author contributions

Yuxing Chen, Q.L. and C.-Z.Z. conceived the project and planned the experiments. Yu Chen and L.W. expressed and purified human ABCC4. Yu Chen and Z.-H.Z. performed cryo-EM data collection, structure determination and model refinement. Yu Chen, W.-T.H. and K.X. performed functional assays. Yuxing Chen, Q.L., C.-Z.Z. and Yu Chen wrote the manuscript.

## Data availability

The cryo-EM structures of apo-form ABCC4, U46619-bound ABCC4 and dipyridamole-bound ABCC4 have been deposited at PDB under the codes of xxx, xxx and xxx, respectively. The cryo-EM density maps of three structures have been deposited at the Electron Microscopy Data Bank (EMD-xxx, EMD-xxx and EMD-xxx, respectively).

---

**Table S1.** Cryo-EM data collection, refinement and validation statistics of ABCC4.

|  | Apo-form<br>ABCC4 | U46619-bound<br>ABCC4 | Dipyridamole-bound<br>ABCC4 |
| --- | --- | --- | --- |
| **Data collection and processing** |  |  |  |
| Magnification | 82,000 | 82,000 | 105,000 |
| Voltage (kV) | 300 | 300 | 300 |
| Electron exposure (e<sup>-</sup>/Å<sup>2</sup>) | 60 | 60 | 60 |
| Defocus range (μm) | -1.5 to -2.0 | -1.5 to -2.0 | -1.5 to -2.0 |
| Pixel size (Å) | 1.06 | 1.06 | 0.82 |
| Symmetry imposed | C1 | C1 | C1 |
| Initial particle images (no.) | 3,407,933 | 2,213,130 | 2,722,564 |
| Final particle images (no.) | 392,047 | 247,335 | 397,218 |
| Map resolution (Å) | 3.13 | 3.08 | 3.4 |
| FSC threshold | 0.143 | 0.143 | 0.143 |
| Map resolution range (Å) | 2.12-999 | 2.12-999 | 1.64-999 |
| **Refinement** |  |  |  |
| Initial model used (PDB code) | XXX | XXX | XXX |
| Map sharpening B factor (Å<sup>2</sup>) | -145.3 | -124.7 | -154.3 |
| Model resolution (Å) | 3.13 | 3.08 | 3.4 |
| FSC threshold | 0.143 | 0.143 | 0.143 |
| **Model composition** |  |  |  |
| Non-hydrogen atoms | 9,784 | 9,809 | 9,820 |
| Protein residues | 1,227 | 1,227 | 1,227 |
| Ligands | 0 | 1 | 1 |
| **R.M.S. deviations** |  |  |  |
| Bond lengths (Å) | 0.01 | 0.022 | 0.015 |
| Bond angles (°) | 1.24 | 1.623 | 1.4 |
| **Validation** |  |  |  |
| MolProbity score | 2.23 | 2.49 | 2.38 |
| Clashscore | 15.57 | 26.99 | 19.33 |
| Rotamers outliers (%) | 0 | 0 | 0 |
| **Ramachandran plot** |  |  |  |
| Favored (%) | 90.52 | 89.35 | 88.19 |
| Allowed (%) | 9.48 | 10.48 | 11.73 |
| Disallowed (%) | 0 | 0.16 | 0.08 |

