## supplementary figures for "Structural insights into human ABCC4 mediated transport of platelet agonist and inhibitor"

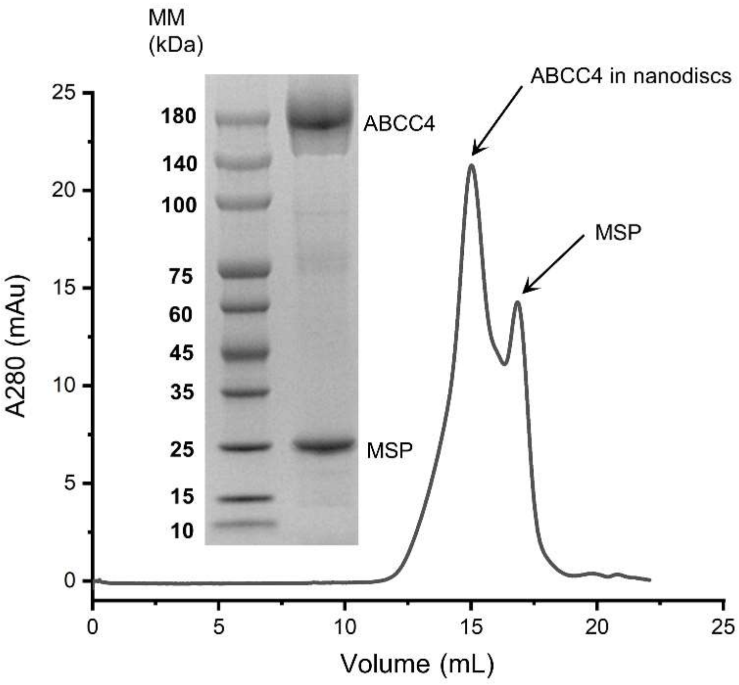


**Extended Data Fig. 1 | Gel filtration profile of purified human ABCC4 in lipid nanodiscs.** The peak fractions were applied to SDS-PAGE and visualized by Coomassie blue staining.

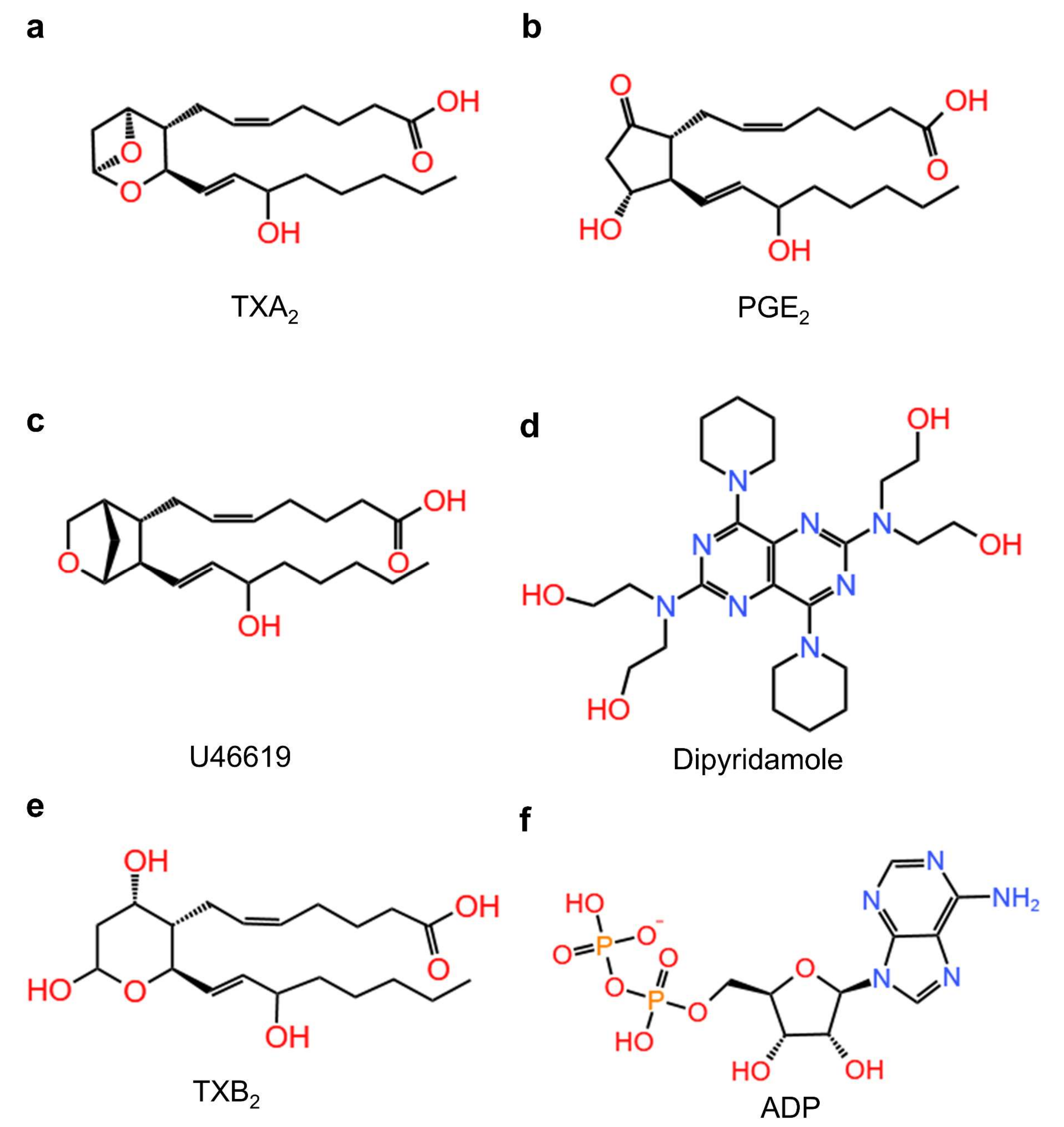


**Extended Data Fig. 2 | Structural formulas of some substrates or inhibitor of ABCC4.**

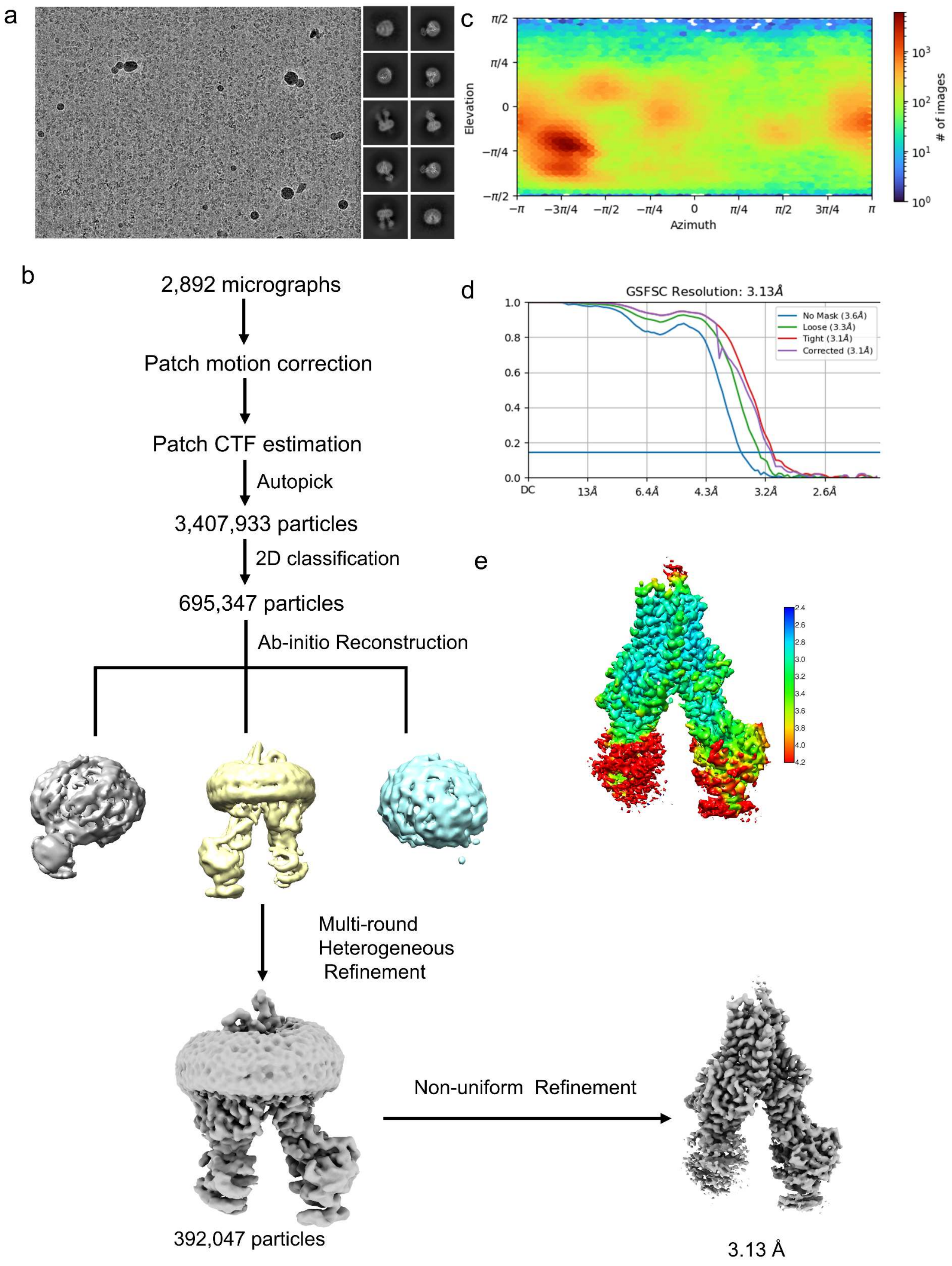
a
c
π/2
π/4
0
−π/4
−π/2
Elevation
−π
−3π/4
−π/2
−π/4
0
π/4
π/2
3π/4
π
Azimuth
### of images
b
2,892 micrographs
Patch motion correction
Patch CTF estimation
Autopick
3,407,933 particles
2D classification
695,347 particles
Ab-initio Reconstruction
Multi-round
Heterogeneous
Refinement
392,047 particles
Non-uniform Refinement
3.13 Å
d
GSFSC Resolution: 3.13Å
No Mask (3.6Å)
Loose (3.3Å)
Tight (3.1Å)
Corrected (3.1Å)
1.0
0.8
0.6
0.4
0.2
0.0
DC
13Å
6.4Å
4.3Å
3.2Å
2.6Å
e
2.4
2.6
2.8
3.0
3.2
3.4
3.6
3.8
4.0
4.2

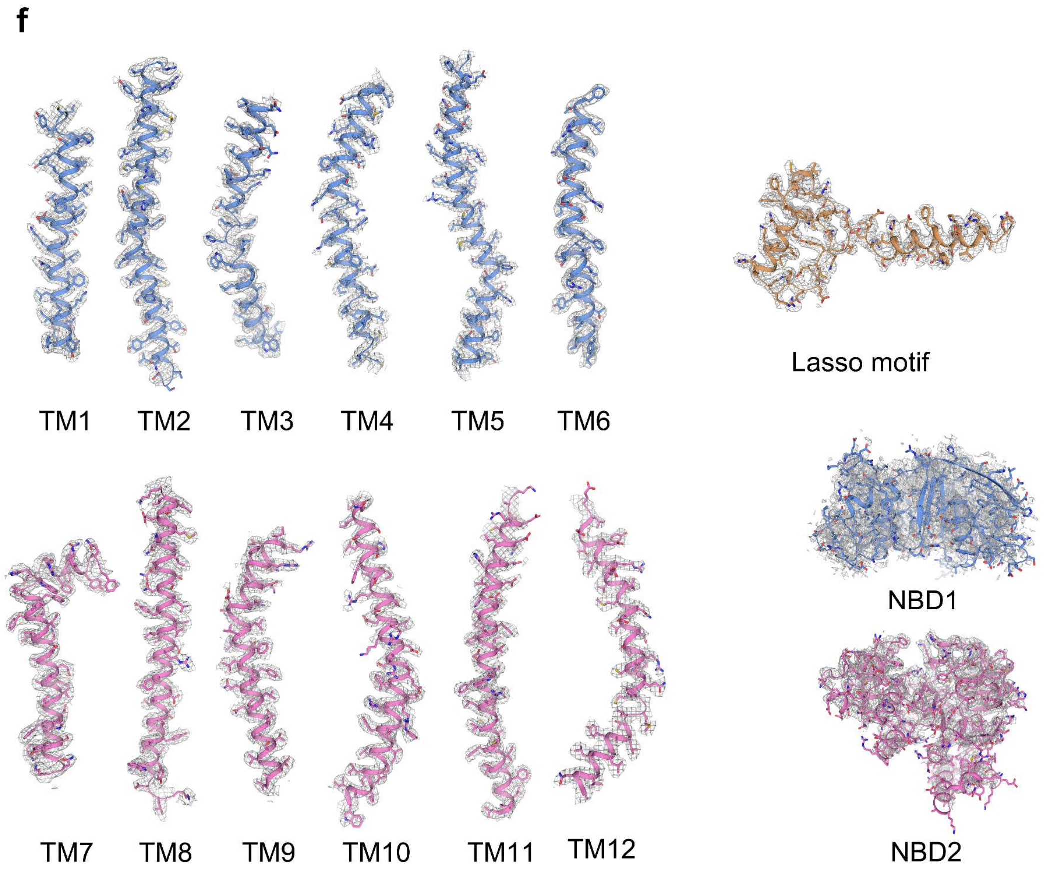


**Extended Data Fig. 3 | Cryo-EM analysis of apo-form ABCC4. a** Representative micrograph and 2D class averages. Bar: 50 nm. The micrograph is a representative of 2892 cryo-EM images. **b** The flowchart for cryo-EM data processing. **c** The angular distribution plot of the final 3-D reconstruction of the apo-form ABCC4. **d** Fourier shell correlation (FSC) curves for the apo-form ABCC4. **e** The local resolution map of apo-form ABCC4. The color code for resolutions, shown with the unit Å, is calculated using CryoSPARC. **f** Cryo-EM maps for representative segments of apo-form ABCC4. The contour level for TM1-12 and the lasso motif are set at 5σ, whereas that for the two NBDs are 3σ.

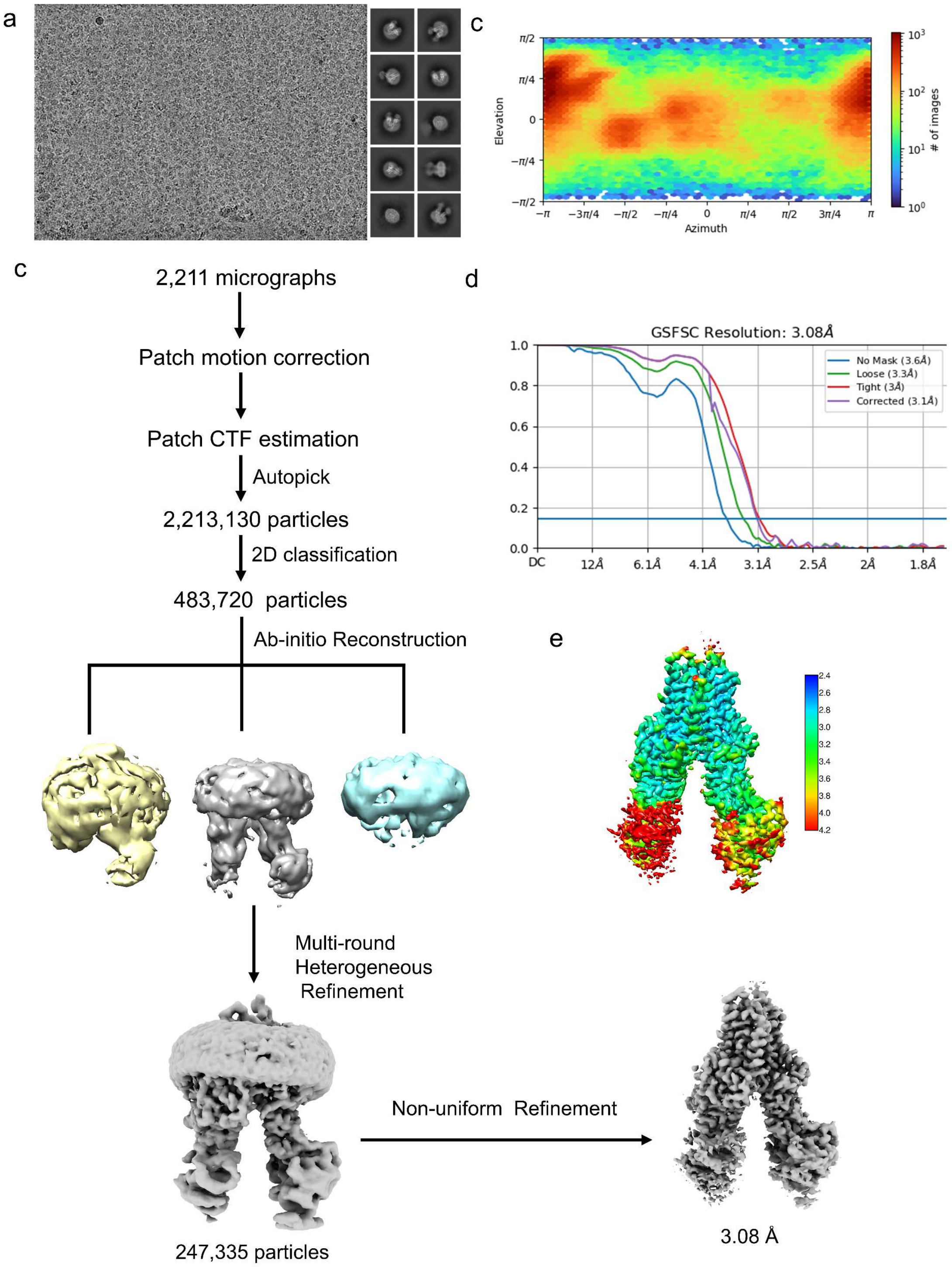
a
c
π/2
π/4
0
−π/4
−π/2
Elevation
−π
−3π/4
−π/2
−π/4
0
π/4
π/2
3π/4
π
Azimuth
### of images
c
2,211 micrographs
Patch motion correction
Patch CTF estimation
Autopick
2,213,130 particles
2D classification
483,720 particles
Ab-initio Reconstruction
Multi-round
Heterogeneous
Refinement
247,335 particles
Non-uniform Refinement
3.08 Å
d
GSFSC Resolution: 3.08Å
No Mask (3.6Å)
Loose (3.3Å)
Tight (3Å)
Corrected (3.1Å)
1.0
0.8
0.6
0.4
0.2
0.0
DC
12Å
6.1Å
4.1Å
3.1Å
2.5Å
2Å
1.8Å
e
2.4
2.6
2.8
3.0
3.2
3.4
3.6
3.8
4.0
4.2

f

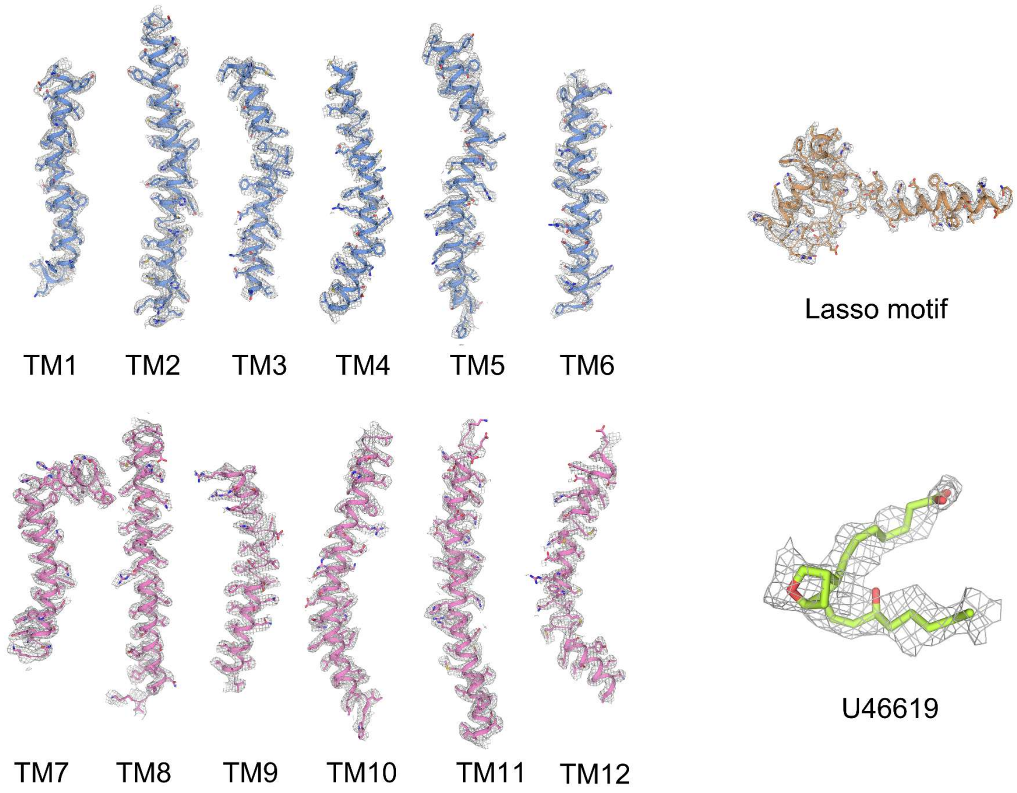


**Extended Data Fig. 4 | Cryo-EM analysis of U46619-bound ABCC4. a** Representative micrograph and 2D class averages. Bar: 50 nm. The micrograph is a representative of 2211 cryo-EM images. **b** The flowchart for cryo-EM data processing. **c** The angular distribution plot of the final 3-D reconstruction of the U46619-bound ABCC4. **d** Fourier shell correlation (FSC) curves for the U46619-bound ABCC4. **e** The local resolution map of U46619-bound ABCC4. The color code for resolutions, shown with the unit Å, is calculated using CryoSPARC. **f** Cryo-EM maps for representative segments of U46619-bound ABCC4. The contour level for TM1-12 and the lasso motif are set at 5σ, whereas that for the U46619 molecule is 3σ.

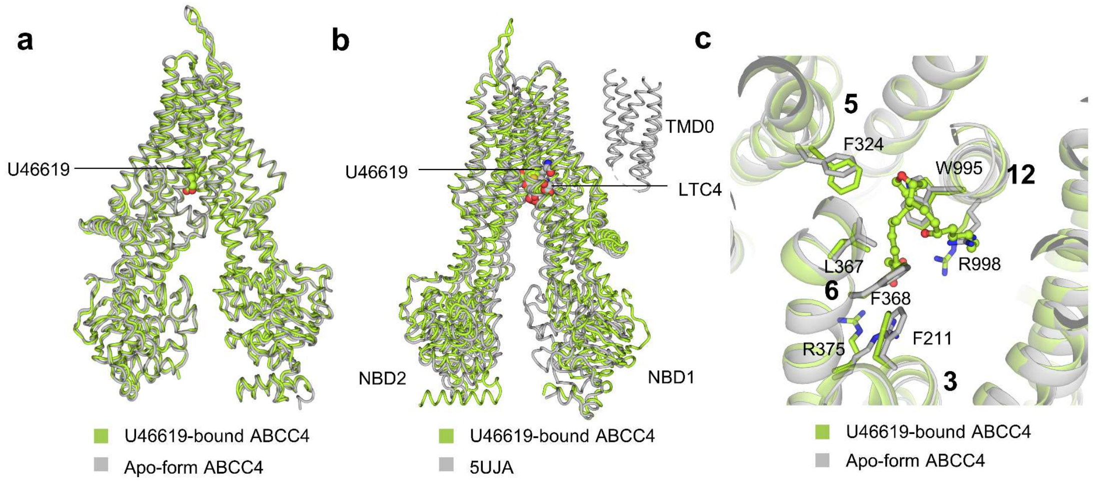


**Extended Data Fig. 5 | Structural comparisons. a** Superposition of U46619-bound ABCC4 (limon) against apo-form ABCC4 (gray). **b** Superposition of U46619-bound ABCC4 (limon) against $LTC_4$-bound bMRP1 (gray, PDB code:5UJA). **c** Superposition of the key residues in the U46619-binding sties (limon) against those of the apo-form ABCC4 (gray). The binding residues are shown as sticks.

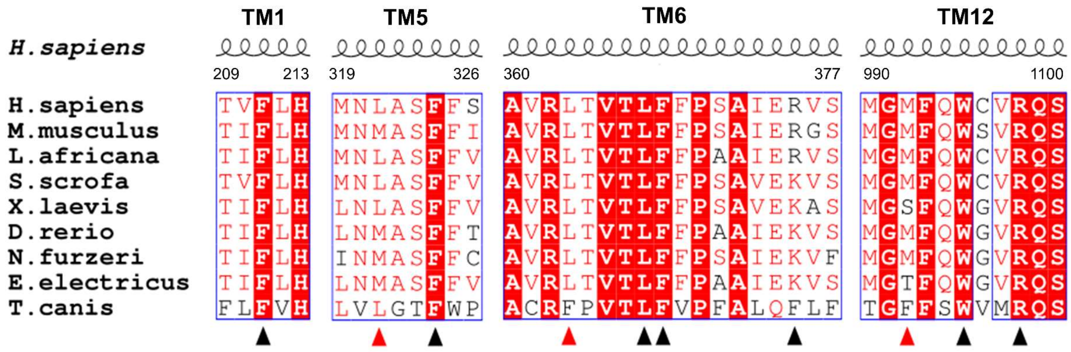


**Extended Data Fig. 6 | Multiple-sequence alignment of ABCC4 and homologs.** Triangles indicate the residues interacting with both substrate and inhibitor (black) or only interacting with dipyridamole (red).

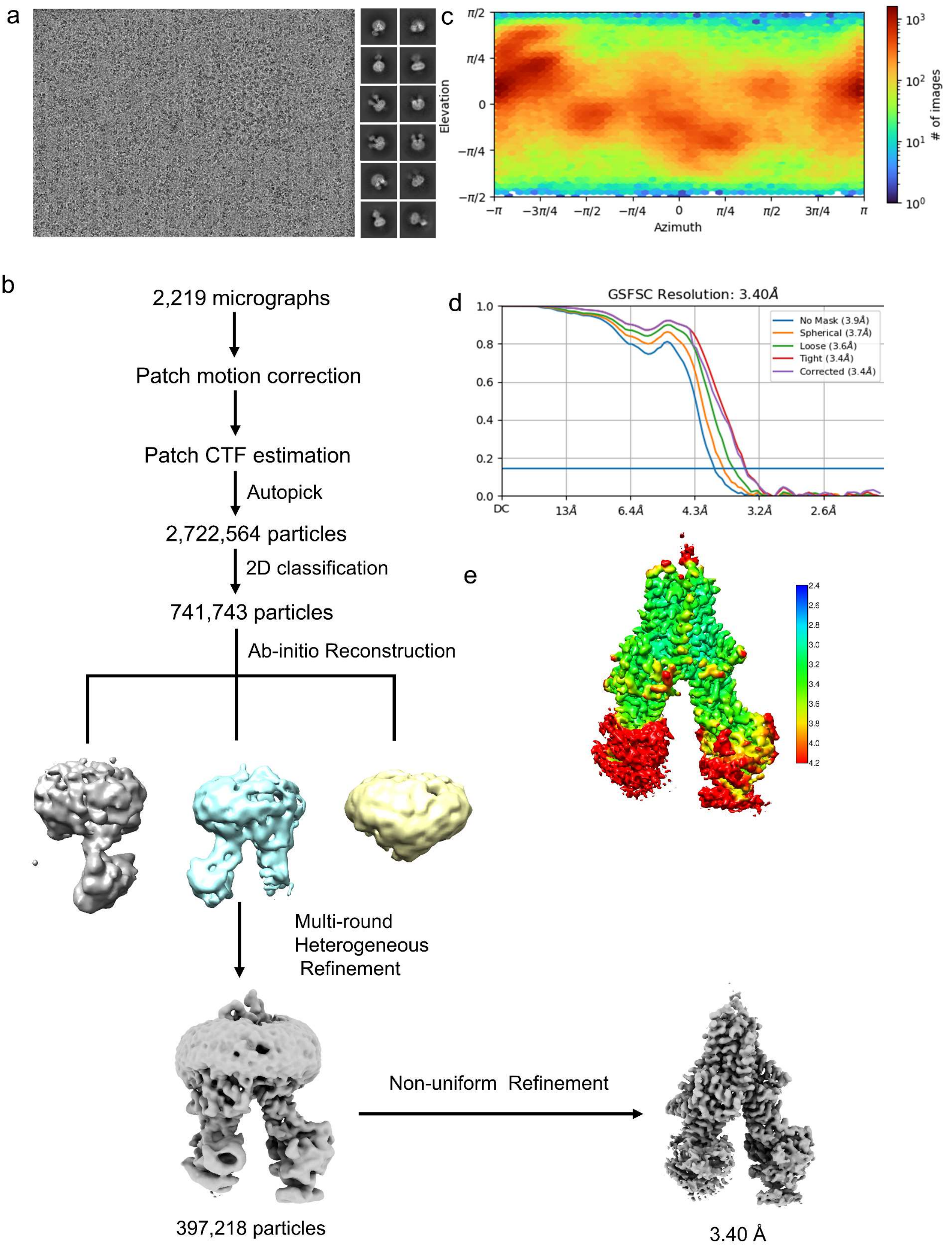
a
c
π/2
π/4
0
−π/4
−π/2
Elevation
−π
−3π/4
−π/2
−π/4
0
π/4
π/2
3π/4
π
Azimuth
### of images
b
2,219 micrographs
Patch motion correction
Patch CTF estimation
Autopick
2,722,564 particles
2D classification
741,743 particles
Ab-initio Reconstruction
Multi-round
Heterogeneous
Refinement
Non-uniform Refinement
397,218 particles
3.40 Å
d
GSFSC Resolution: 3.40Å
No Mask (3.9Å)
Spherical (3.7Å)
Loose (3.6Å)
Tight (3.4Å)
Corrected (3.4Å)
DC
13Å
6.4Å
4.3Å
3.2Å
2.6Å
e

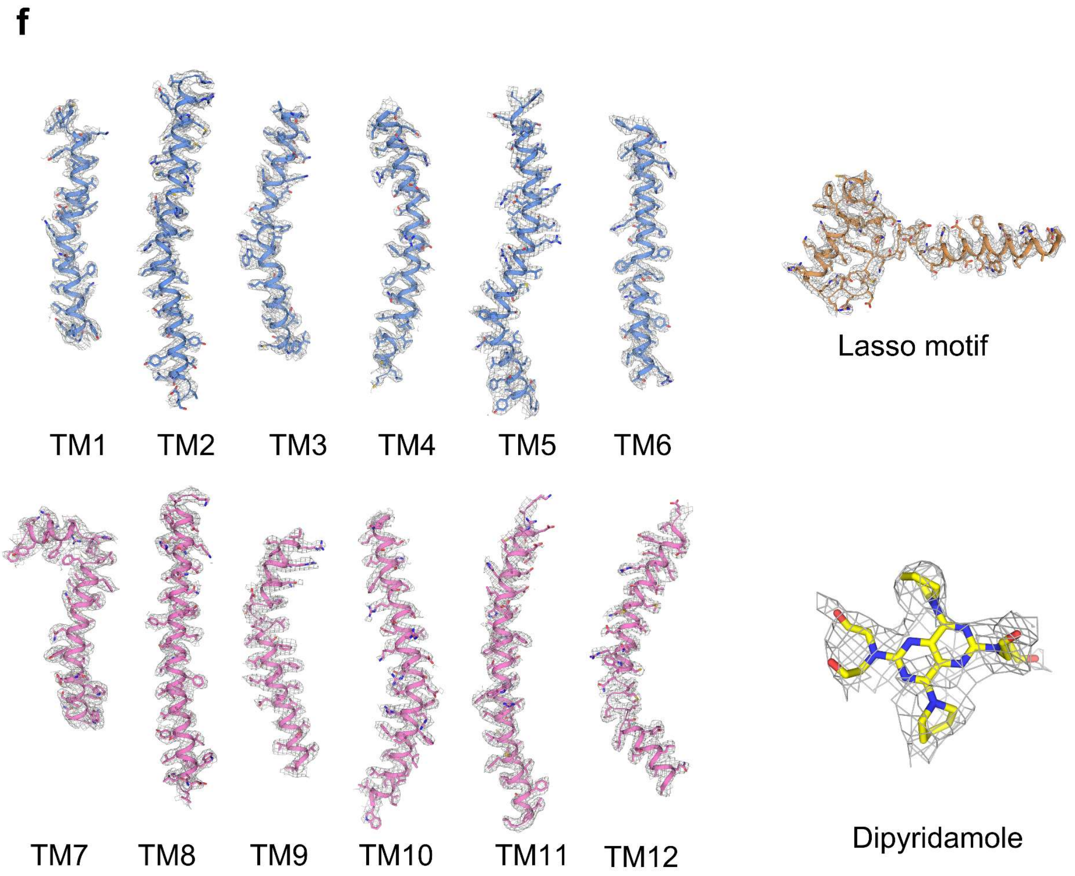


**Extended Data Fig. 7 | Cryo-EM analysis of dipyridamole-bound ABCC4. a** Representative micrograph and 2D class averages. Bar: 50 nm. The micrograph is a representative of 2219 cryo-EM images. **b** The flowchart for cryo-EM data processing. **c** The angular distribution plot of the final 3-D reconstruction of the dipyridamole-bound ABCC4. **d** Fourier shell correlation (FSC) curves for the dipyridamole-bound ABCC4. **e** The local resolution map of dipyridamole-bound ABCC4. The color code for resolutions, shown with the unit Å, is calculated using CryoSPARC. **f** Cryo-EM maps for representative segments of dipyridamole-bound ABCC4. The contour level for TM1-12 and the lasso motif are set at 5σ, whereas that for the dipyridamole molecule is 4σ.

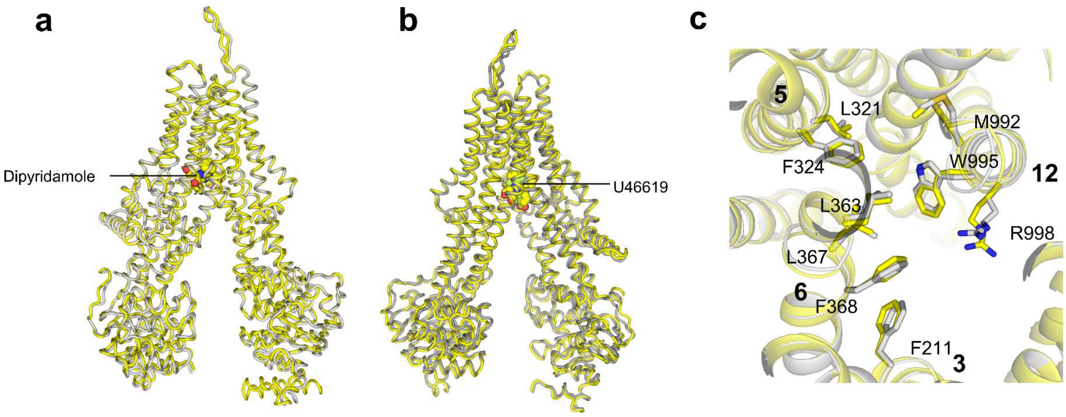


**Extended Data Fig. 8 | Structural comparisons.** Superposition of dipyridamole-bound ABCC4 (yellow) against **a** apo-form ABCC4 (gray) and **b** U46619-bound ABCC4 (limon). **c** Superposition of the key residues in the dipyridamole-binding sties against those of the apo-form ABCC4. The interacting residues are shown as sticks.
